# Protein Docking using Constrained Self-adaptive Differential Evolution Algorithm

**DOI:** 10.1101/312801

**Authors:** S. Sudha, S. Baskar, S. Krishnaswamy

## Abstract

The objective of protein docking is to achieve a relative orientation and an optimized conformation between two proteins that results in a stable structure with the minimized potential energy. Constrained Self-adaptive Differential Evolution (Cons_SaDE) algorithm is used to find the minimum energy conformation using proposed constraints such as boundary surface complementary interactions, non-bonded inter-atomic allowed distances, and finding of interaction and non-interaction sites. With these constraints, Cons_SaDE is efficient enough to explore the promising solutions by gradually self-adapting the strategies and parameters learnt from their previous experiences. Modified sampling scheme called Rotate Only Representation is used to represent a docking conformation. GROMOS53A6 force field is used to find the potential energy. To test the performance of this algorithm, few bound and unbound complexes from Protein Data Bank (PDB) and few easy, medium and difficult complexes from Zlab benchmark4.0 are used. Buried Surface Area, Root Mean Square Deviation (RMSD) and Correlation Coefficient are some of the metrics applied to evaluate the best docked conformations. RMSD values of the best docked conformations obtained from five popular docking web servers are compared with Cons_SaDE results and nonparametric statistical tests for multiple comparisons with control method are implemented to show the performance of this algorithm. Cons_SaDE has produced good quality solutions for the most of the data sets considered.

## 1. Introduction

Recently, drugs which bind to a target protein and then inhibit protein-protein interaction or enzyme reactions have become a subject of interest (Suenaga et al. 2012). The importance of protein interactions have been recently highlighted by large-scale protein-protein interaction maps and have been revealed for many organisms (Li and Kihara 2012). However, it is often difficult to obtain protein complexes by experimental methods. Protein-protein interaction (process called Protein docking) requires efficient sampling across the entire range of positional, conformational and orientation possibilities and hence, this problem is considered as a difficult optimization problem.

From the literature, it is observed that the Fast Fourier Transform (FFT) based algorithms perform an exhaustive global search to find the optimal docking structure and the number of computations used by them is very huge. GRAMM-X (Tovchigrechko and Vakser 2005), ClusPro (Kozakov et al. 2006), ZDOCK (Chen et al. 2003), F2DOCK (Bajaj et al. 2011), DOT (Roberts et al. 2013), ASPDock (Li et al. 2011) and FTDock (Gabb et al. 1997) are few FFT-based docking programs. Disadvantages of this method are that the translational FFT correlations must be repeated for many thousands of rotational samples for one of the molecules. Thus, these docking algorithms are inherently computationally expensive (Banting et al. 2012). HEX (Ritchie et al. 2008) and FRODOCK (Garzon et al. 2009) use spherical functions in Fourier transform, in order to accelerate the search over 3D rotational space. But, when the size of the molecules is huge, these FFT techniques take a longer time to evaluate the optimal pose of binding, since they have to explore a rather large energy landscape.

In local shape geometric hashing methods, local information about shape descriptors is used to find the matches; the algorithms are faster and are able to produce near optimal docking structures. The distance geometry algorithm is used in DOCK (Kuntz et al. 1982), which performs a local search around known binding sites. Geometric hashing is used in PatchDock and SymmDock (Schneidman-Duhovny et al. 2005) and LZerD (Esquivel-Rodrigue et al. 2012) in order to carry out global protein–protein docking search by finding the local matches of shape descriptors. The advantage of these algorithms is their speed and can quickly scan through several thousand ligands in a matter of seconds and figure out the protein’s active site. But a potential disadvantage is the possibility of losing the correct transformation, because of not performing the global search.

In randomized search techniques, the docking structures are randomly generated and are iterated to obtain the minimum energy conformation using the strategies adopted by various algorithms. Since these algorithms are iterative in nature and are able to fine tune the values of the variables, the generation of more accurate docking structures is possible. Examples of randomized search include, Monte Carlo search in RosettaDock (Gray et al. 2003), ATTRACT (de Vries and Zacharias 2013) and pseudo-Brownian Monte Carlo minimization in ICM-DISCO (Fernandez-Recio et al. 2003), Simulated Annealing in HADDOCK (de Vries et al. 2010) and GOLD (Jones et al. 1997), AutoDock (Morris et al. 1998), DIVALI (Clark 1995), MolDock (Thomsen and Christensen 2006) and Lead finder (Stroganov et al. 2008) used evolutionary algorithms, Tabu search methods in PRO LEADS (Baxter et al. 1998) and PSI-DOCK (Pei et al. 2006) and Swarm optimization algorithms in SwarmDock (Moal and Bates 2010), Tribe-PSO (Chen et al. 2006) and PLANTS (Korb et al. 2006). A disadvantage of this method is that the number of computations used for finding the minimum energy conformation is more, because of its iterative process. But, an introduction of constraints into the docking problem make these algorithms have focused search with reduced amount of computations. There are many successful constrained evolutionary algorithms for solving problems with steering search (Michalewicz and Schoenauer 1996; Reid 1996; Coello and Montes 2002; Kong et al. 2013; Cai et al. 2013). So, the design of new perspective of the docking problem with constraints is necessary to have guided search to obtain a group of promising solutions.

For a good docking process, the efficient design of sampling scheme, energy scoring, and the search procedure are necessary. One of the most difficult tasks in computational docking is selecting the best docking conformation from the large search space with reduced number of computations. The proposed approach addresses the new sampling scheme and search procedure. Here, the docking problem is formulated as a constrained optimization problem to produce energy minimized docked conformation satisfying the constraints such as boundary interaction hits, inter-atomic distances and finding of interaction and non-interaction sites. These constraints are introduced into Self-adaptive Differential Evolutionary algorithm, called as Cons_SaDE, to avoid generation of feeble docking structures. To validate the performance of the proposed algorithm, few bound and unbound docking complexes from Protein Data Bank and Zlab Benchmark 4.0 (Zlab 2010) are used.

The flow of this paper is: Section 2 describes the methodology adopted to find the optimized docked conformation; Section 3 discusses the results and finally Section 4 concludes the performance of the algorithm.

## 2. Methodology

In this docking procedure, a new sampling (representation) scheme called Rotate Only Representation (ROR) is proposed for generating promising minimum energy docking conformations. The force field GROMOS 53A6 (Oostenbrink et al. 2004), that considers the strength of physical and chemical forces for scoring, is used for finding the potential energy of those conformations. Three constraints are introduced into this docking problem, namely boundary surface complementary interactions, non-bonded inter-atomic allowed distances and finding of interaction and non-interaction sites, to make Self-adaptive Differential Evolution (SaDE) algorithm as a constrained search algorithm (Cons_SaDE) to have a guided exploration of search space to produce high quality solutions.

### 2.1 Representation

Basically, in rigid docking (Huang et al. 2005), the protein1 is fixed and the protein2 has six degrees of freedom of translation and rotation. Here, Protein2 is allowed to randomly move over protein1 to explore the best docking by having 3 Cartesian coordinates for translation (*Tx, Ty and Tz*), and 3 angles for rotation (*Rx, Ry and Rz*) along X, Y and Z axis in order (3T3R) as in Figure 1(a).

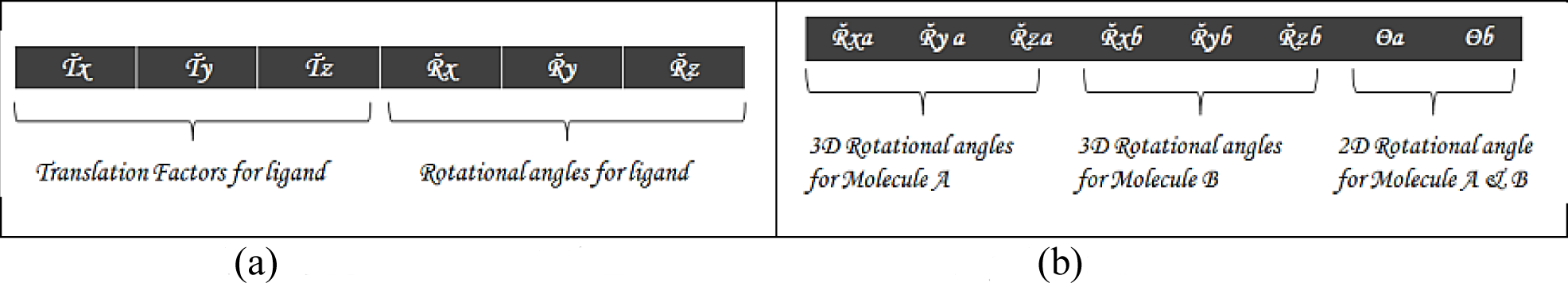
Fig.1(a): 3T3R representation Fig.1(b): ROR representation

In order to improve the search efficiency and to generate more promising solutions, a simple modification is introduced into this sampling scheme. In the proposed Rotate Only Representation (ROR) scheme, both protein1 and protein2 are allowed to rotate. In this scheme, the solutions are represented as a set of randomly generated 8 rotational angles as in Figure 1(b). The locations and orientations of the proteins are random and are not known. The degree of rotation is 360° in either clockwise or anti-clockwise direction.

Of the 8 angles, first 3 angles *(Rxa, Rya, and Rza*) are used for the rotation of protein1 and the next 3 angles (*Rxb, Ryb and Rzb*) for the rotation of protein2. They are rotated along three mutually orthogonal directions, say X, Y, Z. The order of rotations is made on X followed by Y and followed by Z. After rotation, the boundaries of the proteins are derived from their 3D representation and are rotated by *θa* and *θb* about the perpendicular axis to the 2D plane for interaction. Figure 3 depicts the pictorial flow of representation. The rotated boundary surfaces are used for Boundary Surface Complementary Interaction, i.e. to find the shape matching between two proteins using shape complementary property. Here, both proteins are allowed to freely rotate in any direction. This leads to explore all possible combinations of shape matching between proteins.

**Fig.3:**
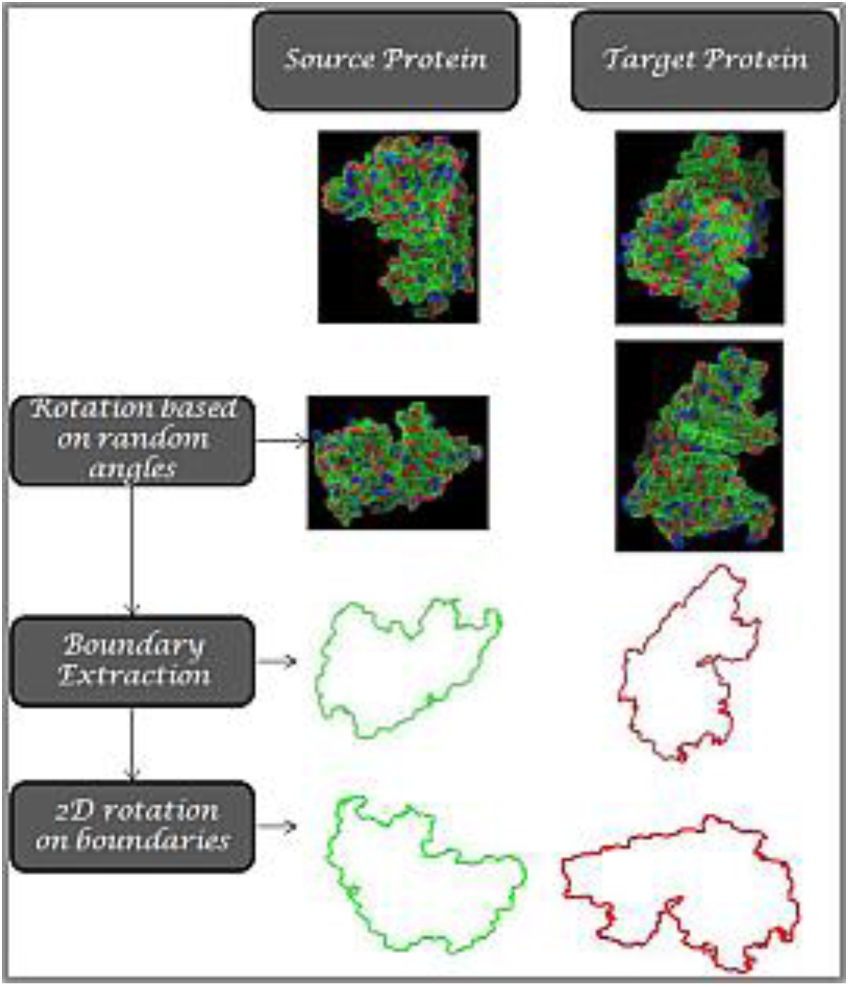
Pictorial flow of representation

To prove the effectiveness of this representation, 80 randomly generated docked solutions adopting ROR sampling scheme and 3T3R scheme in separate are taken. The potential energy of those solutions for each scheme is calculated separately. Good docking solutions will have low potential energy. Figures 4(a)–4(c) show the comparisons of number of minimum energy solutions generated by ROR (blue) and 3T3R (red) sampling schemes. Figure 4(a) shows the comparison results for the data 2SIC, which is an easy complex from Zlab, Figure 4(b) shows the result for 1IJK, a medium complex, whereas Figure 4(c) shows the result of 2OT3, a difficult complex from Zlab. In all three cases, the number of minimum energy solutions generated by ROR is more compared to 3T3R. This is because, in ROR both proteins are freely allowed to rotate in any direction, it is possible to explore more shape matching/ good docking solutions. From these samples, it is known that irrespective of the complex nature of the datasets, ROR scheme generates a large number of minimum energy solutions. Also, ROR never depend on the size of the proteins, whereas in 3T3R, size of the proteins is matter.

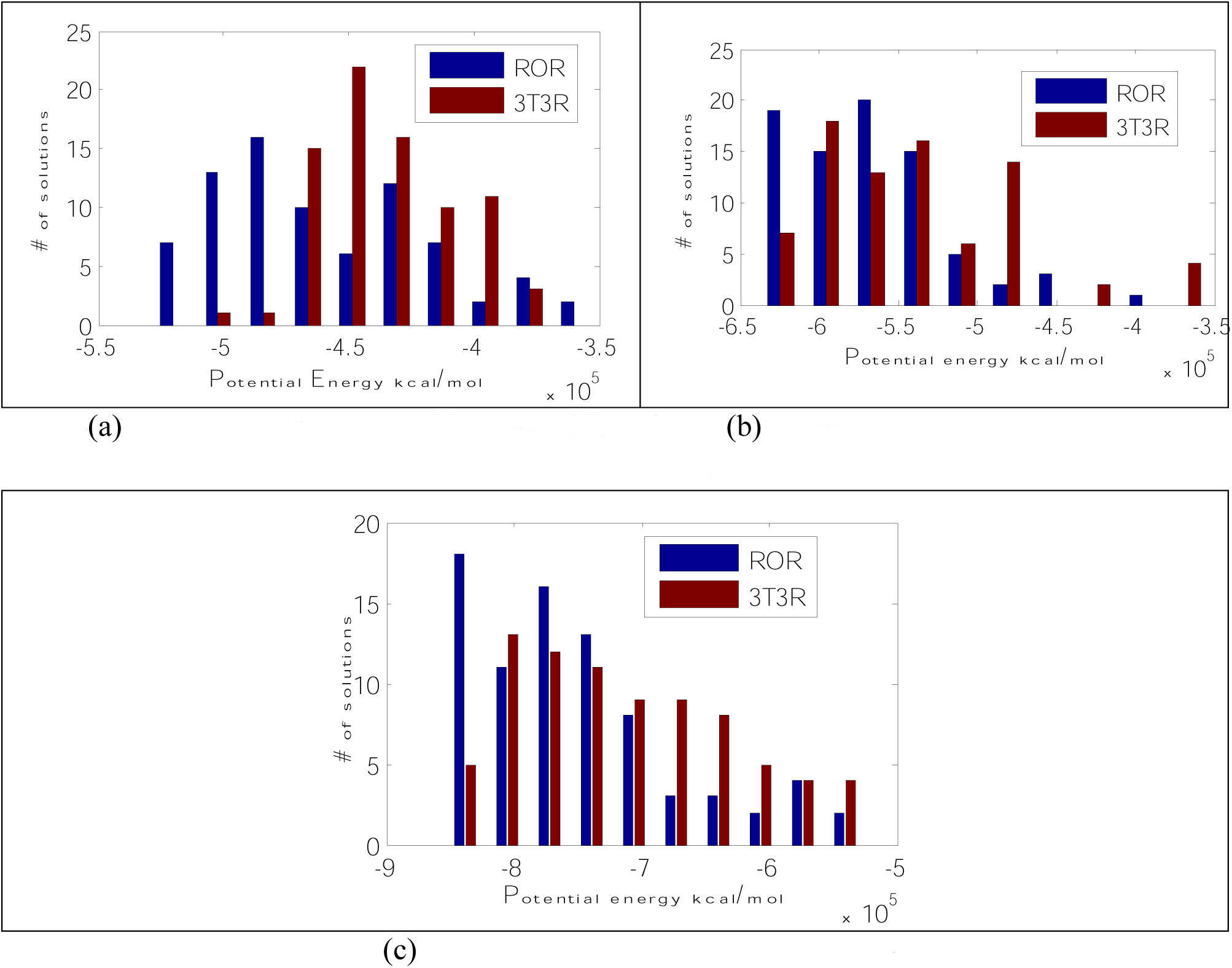
Fig. 4(a): randomly generated solutions for 2SIC Fig. 4(b): randomly generated solutions for 1IJK Fig. 4(c): randomly generated solutions for 2OT3

### 2.2 Fitness Measure

There are several physical and chemical forces involved in the interaction between the two molecules. These forces are used to define various docking scores that measure how good each solution is. These scores take into account the strength of these forces and the plausibility of the docking solution. The force field GROMOS 53A6 (Oostenbrink et al. 2004) is used to find the potential energy of the derived conformations. In the GROMOS force field, the bonded interactions are the sum of bond length, bond angle, harmonic (improper) dihedral angle, and trigonometric (torsional) dihedral angle terms. The nonbonded interactions are the sum of van der Waals (Lennard–Jones, LJ) and electrostatic (Coulomb with Reaction Field, CRF) interactions between all pairs of atoms.

### 2.3 Search Algorithm

The search algorithm generates poses, orientations of particular conformations of the protein in the binding site/ matching part. In order to have fast and guided exploration of the search space to obtain the optimal docked conformation, constrained Self-adaptive Differential Evolution (Cons_SaDE) algorithm is used.

#### 2.3.1 SaDE Algorithm

Differential Evolution (DE) algorithm (Storn and Price 1997) is a powerful population-based stochastic search technique. DE supports several trial vector generation strategies and has three control parameters namely population size(*NP*), scaling factor(*F*), and crossover rate (*CR*) that influence the optimization performance of the DE. The efficiency of DE crucially depends on choosing an appropriate trial vector generation strategy and their associated control parameter values. This is usually done by a trial-and-error scheme which requires high computational costs. The performance of the original DE algorithm is highly dependent on the strategies and parameter settings. Also, during different evolution stages, different strategies with different parameter settings can be more effective than others. Self-adaptation has been found to be highly beneficial for adjusting control parameters during the evolutionary process, especially when done without any user interaction. The Self-adaptive DE (Qin et al. 2009; Cai et al. 2012) algorithm gives a promising path to self-adapt both the trial vector generation strategies and their associated control parameters according to their previous experiences of generating better solutions. The algorithm automatically adapts the trial vector generation strategies and the control parameter *CR* during evolution.

##### Initialization

Randomly generated initial population of size (*NP*) is given by 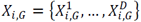, where *i* = 1, … *NP*, *G* = *Generation* and *D* = *Input Dimention*, and it should better cover the entire search space as much as possible by uniformly randomizing individuals within the search space constrained by the prescribed minimum and maximum parameter bounds.

##### Mutation strategy adaptation

The mutation operator is applied to each individual or target vector *X_i,G_* at the generation *G* to produce the mutant vector. After the mutation phase, crossover operation is applied to each pair of the target vector *X_i,G_* and its corresponding mutant vector to generate a trial vector *U_i,G_* Instead of employing the computationally expensive trial-and-error search for the most suitable strategy and its associated parameter values, the SaDE algorithm maintains a strategy candidate pool, which includes four effective trial vector generation strategies with diverse characteristics. For each individual in the current population, one strategy will be chosen according to a probability learned from its previous experience of generating promising solutions; and then applied for the mutation operation. The more successful strategy in the previous generations will get the probability to improve the solutions in the current generation. The strategy candidate pool consists of four strategies as given in Eqs. (1,2,3,4).

- DE/rand/1/bin (ST1)

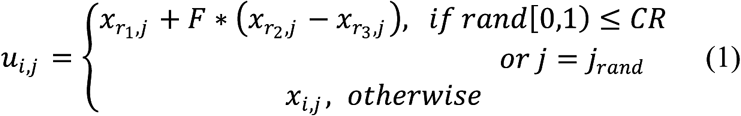
- DE/current-to-best/2/bin (ST2)

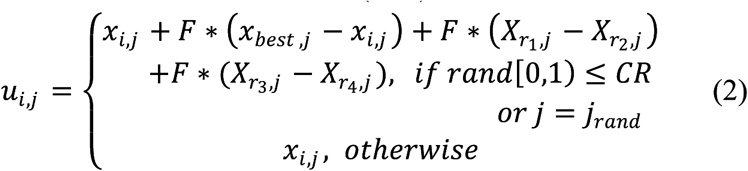
- DE/rand/2/bin (ST3)

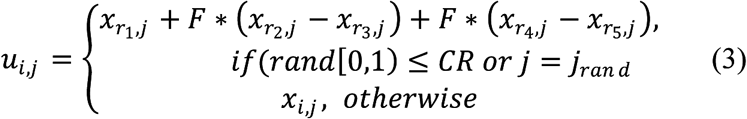
- DE/current-to-rand/1 (ST4)

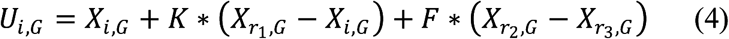

where indices *r*_1_, *r*_2_, *r*_3_ *r*_4_, *r*_5_ are random and mutually different integers generated in the range [1, *NP*], which should also be different from the current trial vector’s index *i*. *F* is a factor in (0, 1+) for scaling the differential vectors and *x_best_,_j_* is the individual vector with the best fitness value in the population. *K* is the coefficient of combination in [-0.5, 1.5].

The binomial-type crossover operator is used in the first three strategies (ST1, ST2 and ST3). The crossover rate *CR* is a user specified constant in the range [0,1). *j_rand_* is a randomly chosen integer in the range [1, D]. The binomial crossover operator copies the *j^th^* parameter of the mutant vector to the corresponding element in the trial vector *U_i,G_*, if *rand* 0,1 ≤ *CR* or *j* = *j_rand_*. Otherwise, it is copied from the corresponding target vector *X_i,j_*. The fourth strategy (ST4) directly generates the trial vector without a crossover. The stochastic universal selection method is used to select one trial vector generation strategy for each target vector in the current population. The probabilities of the strategies are updated only after an initial learning period (*LP*) generation which is set by the user. The probabilities are initialized to 1/*K*; all strategies have equal probability to be chosen. After the initial *LP* generations, the probability of choosing *k^th^* (*k* = 1,2, …, *K*) strategy will be updated at each subsequent generation *p_k,G_* as in Eq.(5),

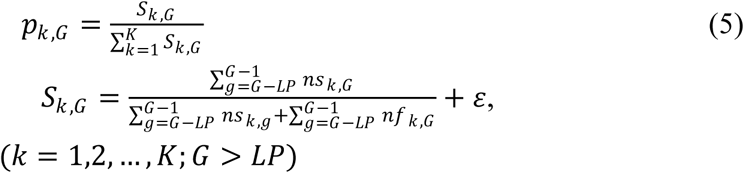

where *S_k,G_* represents the success rate of the trial vectors generated by the *k^th^* strategy and successfully entering the next generation within the previous *LP* generations with respect to generation *G*. The small constant value *ε* = 0.01 is used to avoid the possible null success rates.

##### Parameter Adaptation

The scaling factor *F* is approximated by a normal distribution with mean value 0.5 and standard deviation 0.3. *CR* is approximated as normally distributed variable mean *CRm_k_* with respect to the *k^th^* strategy and standard deviation 0.1. *CRm_k_* is set at 0.5 for all the strategies for an initial *LP* generation. After *LP* generations, the *CRm_k_* value is adapted with the median of the successful *CR* values (that have generated trial vectors successfully entering the next generation) over the past *LP* generations for every subsequent generation.

In the SaDE algorithm, the *F* parameter is approximated by a normal distribution with mean value 0.5 and standard deviation 0.3. A set of values are randomly sampled from such normal distribution and applied to each target vector in the current population. *CR* is normally distributed in a range with mean *CRm_k_* with respect to the *k^th^* strategy and standard deviation 0.1. Initially, *CRm_k_* is set at 0.5 for all the strategies. A set of *CR* values conforming to the normal distribution *N* (*CRm_k_*,0.1) are generated and applied to those target vectors to which the *k^th^* strategy is assigned. To adapt the crossover rate, those *CR* values with respect to the *k^th^* strategy that have generated trial vectors successfully entering the next generation within the previous *LP* generations are stored in *CRMemory_k_*. At each generation after *LP* generations, the median value stored in *CRMemory_k_* will be calculated to overwrite *CRm_k_* Then, values can be generated according to the new *CRm_k_* for the *k^th^* strategy. After evaluating the newly generated trial vectors, *CR* values in *CRMemory_k_* that correspond to earlier generations will be replaced by promising *CR* values obtained at the current generation with respect to the *k^th^* strategy. The control parameter *K* in the strategy “DE/current-to-rand/1” is randomly generated within [0,1] so as to eliminate one additional parameter. Figure 5 shows the flow diagram of SaDE algorithm.

##### Constrained SaDE (Cons_SaDE)

Three constraints are introduced in SaDE to have fast and guided exploration of the search space to obtain the best conformation. These constraints on solutions help to limit the search space and improve the quality of solutions. To handle the constraints, ε–constraint Relaxation scheme (Takahama and Sakai 2009) is used.

**Fig. 5:**
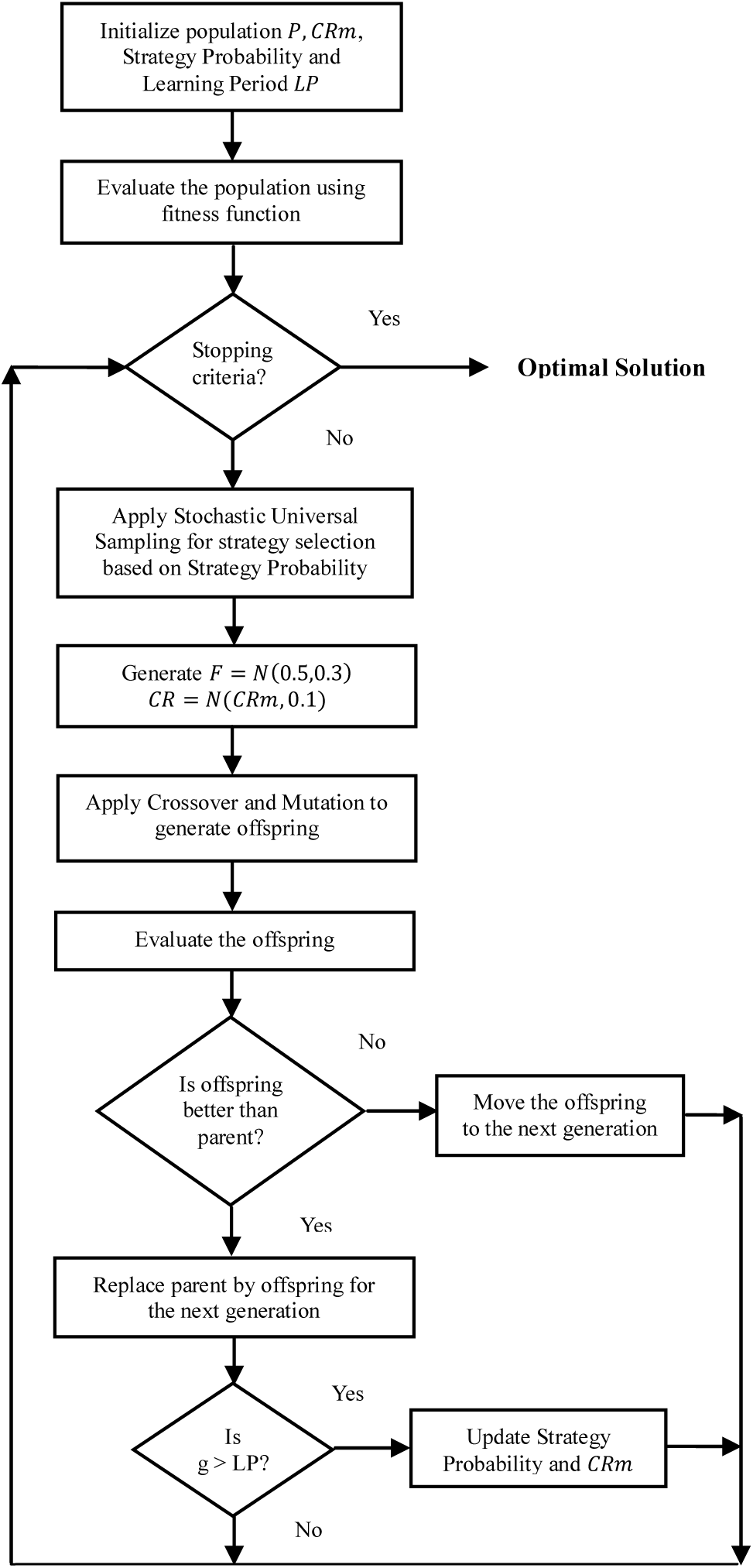
Algorithmic flow of SaDE

The generated conformations (solutions) undergo constraints testing. Solutions satisfying all constraints will be selected for the next generation. Penalty will be added to solutions, which do not satisfy constraints. The amount of penalty added will depend on the amount of deviation of constraints. Solutions satisfying none of these constraints have more probability to be rejected by the algorithm. If a solution fails to satisfy one or two constraints with less violation, Cons_SaDE retunes the parameter values of this solution by learning from its previous experiences and promotes it to a promising solution for the next generation.

The measurements used for validating the constraints are different. They are normalized by scaling between 0 and1 using 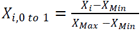, where *X_i_* is each data point *i, X_Min_* is the minimum among all the data points, *X_Max_* is the maximum among all the data points and *X_i,0_ to* 1 is the data point *i* normalized between 0 and 1.

#### 2.3.2 Description of Constraints

##### Boundary Surface Complementary Interaction

In shape complementary methods, the complementary between two surfaces leads to shape matching and helps to find the complementary pose of proteins involve in docking.

Boundary Surface Complementary Interaction procedure is used to calculate the amount of interaction between two proteins’ complementary poses. This is achieved using ROR sampling scheme. ROR scheme introduces randomness through rotational angles over the proteins to recover the interaction pose. After 2D rotations, the boundary of second protein is translated arbitrarily large enough along X axis to counterpart the surface of the first protein. The resulting protein pair geometries are analyzed for shape complementary matching and the amount of interaction is calculated. It is called interaction hits. Figure 6 depicts the flow of this procedure.

**Fig. 6:**
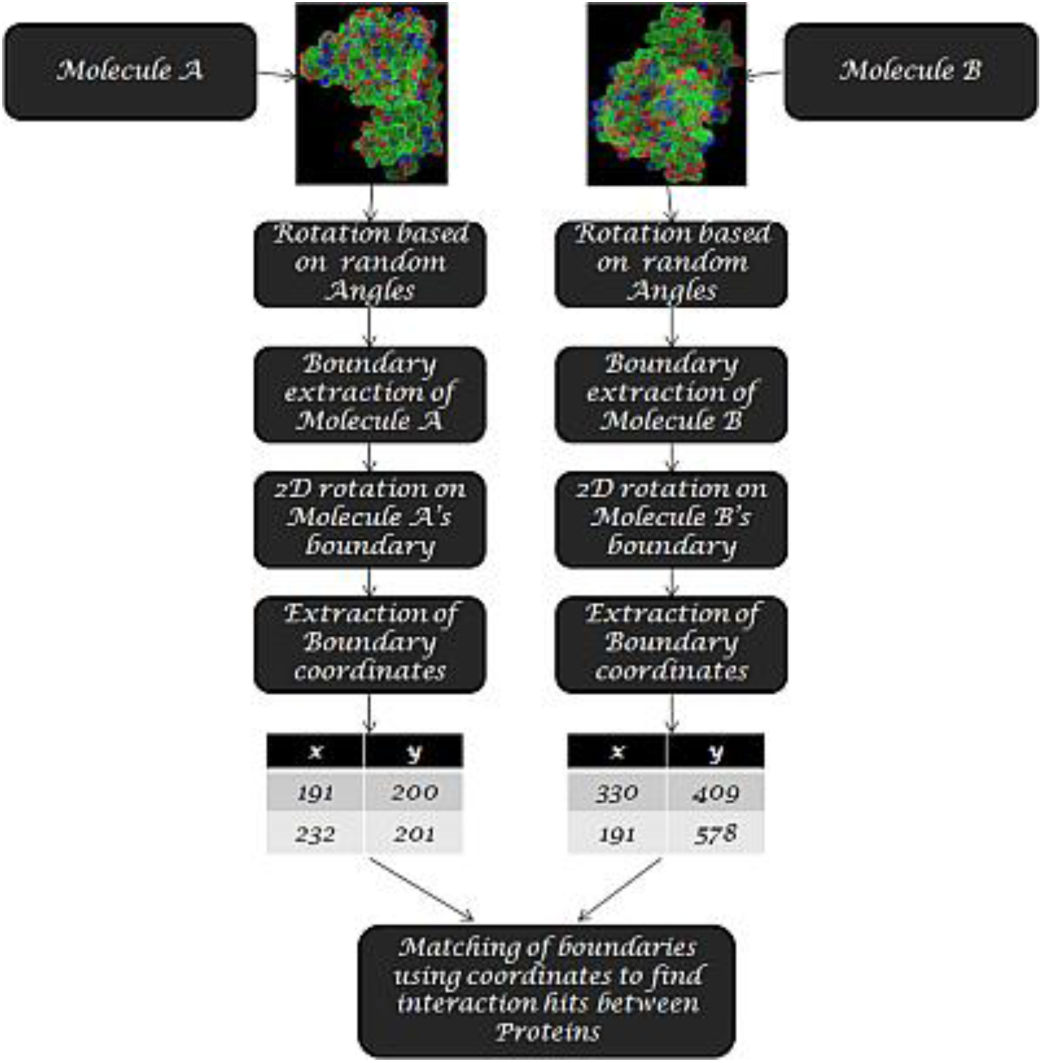
Algorithmic flow of constraint I

Shape complementary approaches are robust and can easily figure out the matching of proteins. So, Boundary Surface Complementary Interaction procedure is used as one of the constraints to find the docking solution having maximum interaction hits. When the generated conformation exhibits lesser interaction hits than the threshold value, it is treated as constraint violated solution. Penalty is added according to its violation and it is handled by Cons_SaDE.

In FFT and geometric hashing approaches, more number of docking poses is required to explore the best docking solution, when the size of proteins increases, whereas ROR does not depend on size of the proteins.

##### Avoiding short contacts

This constraint is used to maintain the minimum distance between non-bonded inter atoms to avoid inter penetration of proteins. The generated conformations are tested for minimum non-bonded allowed inter atomic distances. For a stable conformation, bonded inter atomic distances between any two non-hydrogen atoms should be typical between 2.8 Å and 3.5Å. Solutions with distances less than 2.8Å and greater than 3.5Å are treated as constraint violated. Normalized constraint values between these two ranges (0 to 2.7 Å and greater than 3.5 Å) are calculated and penalty will be added accordingly. Cons_SaDE decides whether to ignore the penalized solution or to retune the parameter values of this solution by learning from its previous experiences and promotes it to a promising solution for the next generation.

##### Finding of interaction and non-interaction sites

The purpose of this constraint is to ensure the retention of a docked conformation for further generations based on its correct interaction and non interaction sites. The interaction and non-interaction sites are identified from Accessible Surface Area (ASA) of each residue. The dssp software (dssp 2012) is used to find the ASA.

Once a docked structure is generated, the ASA of each residue is compared with its respective residue’s ASA of the reference structure. If ASAs of the residues are zero in both docked and reference structures, they are interacting (True Positive-TP) residues, because they are buried. If non-zero in both structures, they are non-interacting residues (True Negative-TN). Mismatches are false positives (FP) and false negatives (FN) accordingly. A good docking structure should have minimum false positives and false negatives. High false positives and false negatives, being detrimental to successful protein-protein interaction, need to be avoided. So, a docking structure with high FN and FP are treated as constraint violated solutions.

Instead of handling TP, TN, FP, and FN directly, the measures sensitivity and specificity are precisely used as constraints here. Sensitivity, the true positive rate, measures the proportion of actual interaction sites which are correctly identified. Specificity, the true negative rate, measures the proportion of actual non-interaction sites which are correctly identified. The sensitivity and specificity of a conformation can be calculated as in Eq.(6).

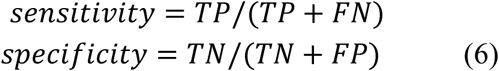

It is decided by trial and error in Cons_SaDE that solutions with more than 75% sensitivity and specificity are good conformations. Other solutions are treated as constraint violated solutions and penalty will be added to the solution. These solutions may be ignored by the algorithm or fine tuned using the crossover and mutation operators (when the number of TP and TN are better) and may be considered for the next generation.

Hence, every solution generated by Cons_SaDE undergoes these three constraints testing i.e. to have a maximum shape complementary matching, avoiding inter-twining of proteins while docking and to ensure maximum interaction and non interaction sites of a docked conformation. A solution satisfying these constraints is easily promoted to next generation as a promising solution. Violated solutions are ignored or reconsidered by Cons_SaDE by learning from its previous experiences.

#### 2.3.3 Fitness Algorithm for Cons_SaDE

The steps given below describe the procedure for fitness calculation.

~~~
***Input***    8 rotational angles between-360° and 360°: 3 for
         protein1’s rotation, 3 for protein2’s rotation and 2 for 2D rotations to represent a solution
***Output***   Potential energy and sum of normalized constraint violations
***Step 1***   Protein1 is loaded in molecular visualization software and it undergoes rotation
         based on randomly generated angles along X, Y and Z directions. Then, Protein2 is
         loaded and rotated based on its random angles. Here, the orientation and location of
         two proteins are random.
***Step 2***   The boundaries of rotated structures are extracted.
***Step 3***   2D rotation is applied on the boundaries of proteins using 2 random angles. The
         docked structure after rotations is saved.
***Step 4***   Boundary coordinates of both rotated proteins are extracted.
***Step 5***   Protein2’s boundary is translated arbitrarily large enough along the X axis by
         connecting the protein1’s surface points so that they coincide.
***Step 6***   Now, using the matched boundary coordinates, interaction hits between two proteins
         are calculated and checked for violation.
***Step 7***   The docked conformation undergoes potential energy calculation using GROMOS     53A6 force field.
***Step 8***   Inter atomic distance between all atoms are calculated for the docked structure and
         checked for violation.
***Step 9***   Sensitivity and specificity measures are calculated and checked for violation.
***Step 10***  Normalized values of interaction hits, inter atomic distances and sensitivity and
         specificity measures, are added. Constraint violations and potential energy are used
         as fitness measures of a solution.
~~~

## 3. Results and Discussion

Cons_SaDE algorithm is executed for 10 independent trials. It explored the search space efficiently and produced multiple conformations with minimum potential energy satisfying the above-mentioned constraints. The best conformation with minimum potential energy is considered for further analysis.

### 3.1 Data Description

It is believed that a newly proposed approach needs first to be tried out in a setting that allows isolating effects and drawing some earlier lessons learnt about the real-world use of the approach. Hence, inline with the previous research (Moal and Bates 2010; Hashmi and Shehu 2012; Esquivel-Rodríguez and Kihara 2012; Mashiach et al. 2010; Chaudhury and Gray 2008; Wang et al. 2007), the proposed algorithm is validated with a small dataset. This dataset consists of few random samples from rigid body, medium difficulty and difficult categories from Zlab benchmark 4.0 and few bound and unbound complexes from Protein Data Bank and is shown in Table 1. Also, use of this dataset is only for illustration purpose of the proposed algorithm.

**Table 1:** Test Data Description

| Protein Complexes | Protein1 | Protein2 | Category |
| --- | --- | --- | --- |
| ***Bound(PDB)*** |  |  |  |
| 4PHV | HIV-1 Protease_A | HIV-1 Protease_B | Inhibitor |
| 1MSB | Mannose-Binding Protein_A | Mannose-Binding Protein_B | Antibody/Antigen |
| 1BMT | Methionine Synthase_A | Methionine Synthase_B | Others |
| ***Unbound(PDB)*** |  |  |  |
| 1CSE | Subtilisin Carlsberg | Eglin C | Enzyme/Inhibitor |
| 2PTC | Beta-Trypsin | Trypsin Inhibitor | proteinase/inhibitor |
| 1ABI | Alpha-Thrombin (Small) | Alpha-Thrombin (Large) | Enzyme/Substrate |
| 4HMG | Hemagglutinin | Hemagglutinin | Others |
| 2MCP | Ga-Kappa Mcpc603 Fab (LC) | Iga-Kappa Mcpc603 Fab (HC) | Others |
| ***Easy(Zlab)*** |  |  |  |
| 2SIC | Subtilisin | Streptomyces subtilisin inhibitor | Enzyme/Inhibitor |
| 1DFJ | Ribonuclease A | Rnase inhibitor | Enzyme/Inhibitor |
| 1BUH | CDK2 kinase | Ckshs1 | Others |
| ***Medium(Zlab)*** |  |  |  |
| 1ACB | Chymotrypsin | Eglin C | Enzyme/Inhibitor |
| 1IB1 | 14-3-3 protein | Serotonin N-acteylase | Others |
| 1IJK | Von Willebrand Factor dom.A1 | Botrocetin | Enzyme/Inhibitor |
| 2NZ8 | Rac GTPase | DH/PH domain of TRIO | Others |
| ***Difficult(Zlab)*** |  |  |  |
| 1JZD | DsbC disulfide bond isomerase | DsbD disulfide bond isomerase | Others |
| 1PXV | Cystein protease | Cystein protease inhibitor | Enzyme/Inhibitor |
| 2C0L | PTS1 and TRP region of PEX5 | SCP2 | Others |
| 2O3B | NucA nuclease | NuiA nuclease inhibitor | Enzyme/Inhibitor |
| 2OT3 | Rab21 GTPase | Rabex-5 VPS9 domain | Others |

### 3.2 Evaluation of Results

The quality of the solutions is analyzed using metrics like buried surface area, RMSD, accuracy and Matthew’s correlation coefficient. As sensitivity and specificity are considered as constraints, the docked structure with minimum energy will have, by design, more than 75% sensitivity and specificity. A good docking structure exhibits minimum potential energy, more buried surface area, and least RMSD values.

#### 3.2.1 Buried Surface Area and RMSD Calculation

According to shape complementary, when shape matching with the complementary pose of two proteins is more, a number of atoms that get buried are also more. Accessible Surface Area for buried atoms is zero. According to first constraint, when the docked structure has more interaction hits, its buried surface area is more.

Buried surface area can be calculated in the following manner. First, Accessible Surface area (ASA) of the docked structure is calculated. Then, protein1 and protein2 are separated from the docked structure and their accessible surface areas are calculated. Then, buried surface area is as in Eq. (7).

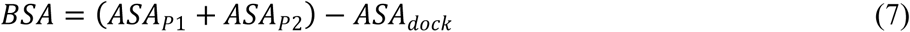

where *P*1 is Protein 1 *P*2 is Protein 2, *dock* is docked structure.

Root-mean-square deviation (RMSD) is the measure of the average distance between the atoms (usually the backbone atoms) of superimposed proteins. Here, RMSD is calculated using “align command with cycles 0” in Pymol (Pymol 2000).

Table 2 shows results of minimum potential energy, buried surface area, iRMSD and ranking of good docked structure of various test data. The potential energies for the docked structures obtained from Cons_SaDE are compared with potential energies of the reference structures and they are found good. Also, the buried area of docked structures is closer to the buried area of their reference structures. interface RMSD (iRMSD) is RMSD value between interacting residues of docked and reference structure. The results show that iRMSD is good and less than 1for all cases, except for medium and difficult categories.

Rank is given based on potential energy and RMSD value of a docked solution. The ranks are assigned to the solutions obtained from the last generation of Cons_SaDE. Out of 20 test data complexes considered here, 19 complexes have ranks less than 100 (i.e. 95% of the test data are having ranks less than 100). Out of these 19 complexes, 11 complexes have ranks less than 10, which is 58%. Among these 11 complexes, 6 complexes are in the first rank (55%). These results show that the docked structures with minimum potential energy possess the least RMSD values.

From the arithmetical results, it can be inferred that the algorithm can produce better docking structures with potential energy, buried surface area, and RMSD values in good ranks.

**Table 2:** Docking Results-Potential Energy, Surface Area, iRMSD and Rank

| Table 2: Docking Results- Potential Energy, Surface Area, iRMSD and Rank |  |  |  |  |  |  |
| --- | --- | --- | --- | --- | --- | --- |
| Data Set | Min. Potential energy |  | Buried Surface Area Å<sup>2</sup> |  | Docked structure |  |
|  | kcal/mol |  |  |  |  |  |
|  | Reference structure | Docked structure | Reference structure | Docked structure | iRMSD (Å) | Rank |
| *Bound(PDB)* |  |  |  |  |  |  |
| 4PHV | -3.09E+05 | -2.91E+05 | 3115.9 | 3114.5 | 0.05 | 1 |
| 1MSB | -5.38E+05 | -5.90E+05 | 1223.2 | 1220.0 | 0.07 | 1 |
| 1BMT | -6.83E+05 | -7.46E+05 | 1530.1 | 1523.0 | 1.97 | 9 |
| *Unbound (PDB)* |  |  |  |  |  |  |
| 1CSE | -3.32E+05 | -4.16E+05 | 9180.2 | 9291.6 | 0.03 | 17 |
| 2PTC | -3.82E+05 | -3.90E+05 | 1451.0 | 1444.3 | 0.17 | 38 |
| 1ABI | -3.44E+05 | -3.45E+05 | 2474.2 | 2297.7 | 0.10 | 107 |
| 4HMG | -2.33E+06 | -3.39E+06 | 21285.2 | 20483.2 | 0.05 | 3 |
| 2MCP | -5.77E+05 | -5.80E+05 | 3690.8 | 3755.2 | 0.10 | 1 |
| *Easy(Zlab)* |  |  |  |  |  |  |
| 2SIC | -4.49E+05 | -4.68E+05 | 1687.5 | 1652.8 | 0.74 | 3 |
| 1DFJ | -6.44E+05 | -6.79E+05 | 2905.2 | 2897.4 | 2.75 | 4 |
| 1BUH | -5.54E+05 | -5.69E+05 | 1367.3 | 1419.7 | 0.08 | 57 |
| *Medium (Zlab)* |  |  |  |  |  |  |
| 1ACB | -3.97E+05 | -3.99E+05 | 1607.1 | 1550.9 | 0.35 | 42 |
| 1IB1 | -9.77E+05 | -9.27E+05 | 3293.0 | 4026.9 | 2.05 | 4 |
| 1IJK | -6.91E+05 | -6.47E+05 | 1378.4 | 1038.6 | 1.10 | 1 |
| 2NZ8 | -6.82E+05 | -6.31E+05 | 1920.0 | 1841.3 | 1.72 | 22 |
| *Difficult(Zlab)* |  |  |  |  |  |  |
| 1JZD | -7.45E+05 | -7.11E+05 | 2138.2 | 874.0 | 5.92 | 25 |
| 1PXV | -4.68E+05 | -4.52E+05 | 2407.1 | 2724.1 | 1.25 | 1 |
| 2C0L | -5.56E+05 | -8.53E+05 | 1783.5 | 2104.3 | 5.14 | 92 |
| 2O3B | -4.79E+05 | -5.87E+05 | 464.6 | 399.2 | 0.58 | 18 |
| 2OT3 | -9.05E+05 | -8.59E+05 | 402.2 | 302.8 | 4.47 | 1 |

Figure 7 shows the best superimpose of docked structure with its reference structure from each category.

**Fig. 7:**
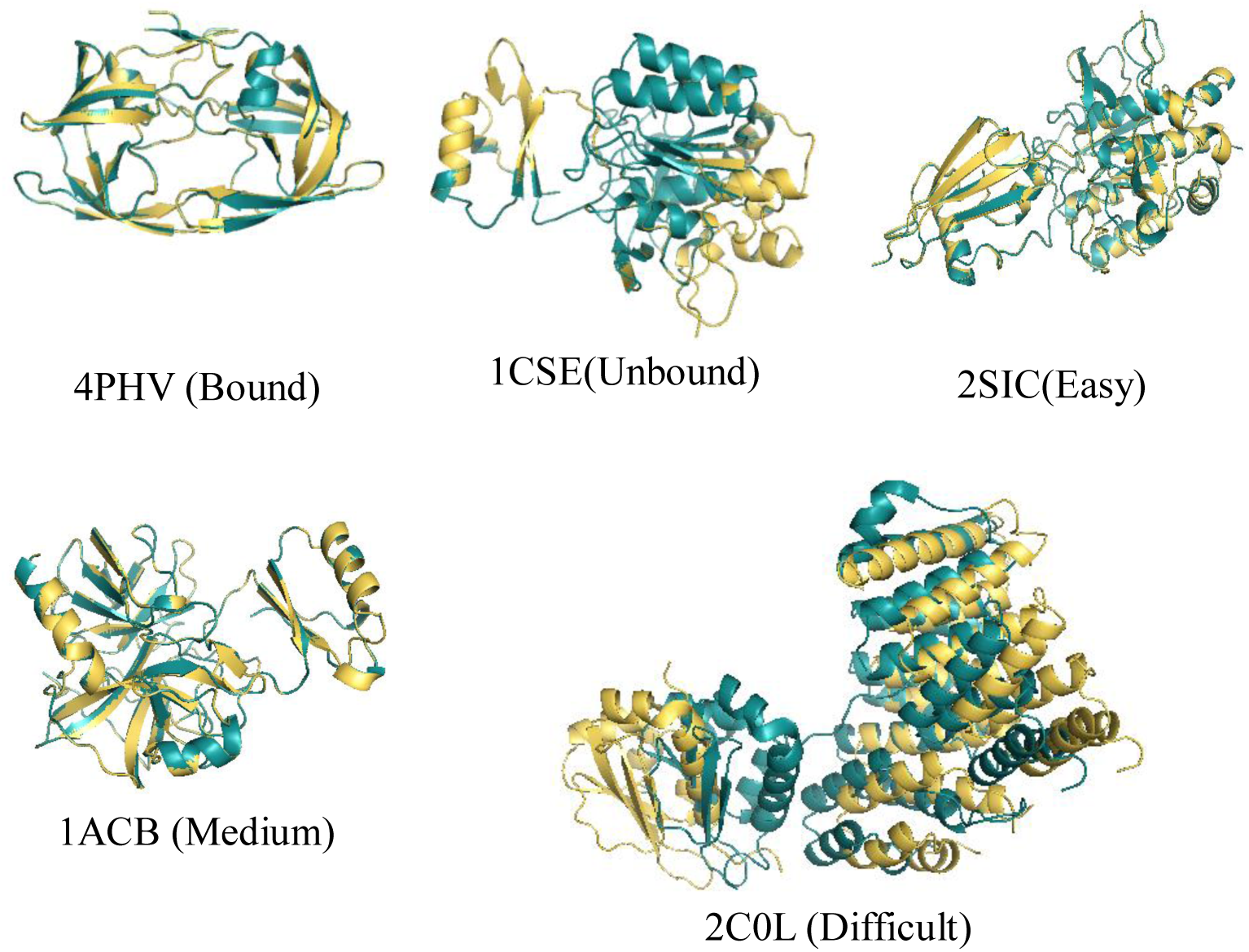
Superimposition of structures (pink-reference structure; green-derived structure)

#### 3.2.2 Accuracy and Correlation Coefficient

To assess the best energy conformations obtained by Cons_SaDE, the statistical measures like sensitivity, specificity, accuracy and Matthew’s Correlation Coefficient (MCC) are used. Sensitivity and specificity are used as constraints in the algorithm. Accuracy and correlation coefficient of a conformation can be calculated using Eq. (8).

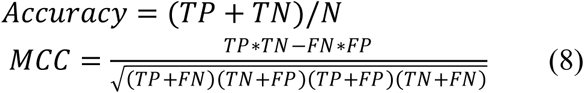

Table 3 shows the results of these measures. Accuracy is a dependent measure and it will become biased towards that highest value of either sensitivity or specificity. From Table 3, it is clear that accuracy depends on specificity. Correlation coefficient reflects the sensitivity values. When sensitivity is less, the correlation coefficient is also less and vice versa. However, MCC often provides a better-balanced evaluation of prediction. MCC returns a value between −1 and +1. A coefficient of +1 represents a perfect prediction, 0 represents no better than random prediction and −1 indicates total disagreement between prediction and observation. A correlation score greater than 0.8 is described as *strong*, whereas less than 0.5 is described as *weak* (Correlation 2016).

**Table 3:** Results of Statistical Measures

| Protein Structures | Sensitivity % | Specificity % | Accuracy % | MCC |
| --- | --- | --- | --- | --- |
| *Bound* |  |  |  |  |
| 4PHV | 83.33 | 98.84 | 96.43 | 0.85 |
| 1MSB | 96.88 | 99.49 | 99.12 | 0.96 |
| 1BMT | 87.27 | 96.78 | 95.71 | 0.80 |
| *Unbound* |  |  |  |  |
| 1CSE | 98.80 | 99.21 | 99.11 | 0.98 |
| 2PTC | 88.24 | 99.16 | 97.80 | 0.90 |
| 1ABI | 92.86 | 98.33 | 90.70 | 0.90 |
| 4HMG | 95.24 | 98.41 | 97.62 | 0.94 |
| 2MCP | 96.36 | 99.22 | 98.86 | 0.95 |
| *Easy* |  |  |  |  |
| 2SIC | 94.87 | 98.34 | 97.63 | 0.93 |
| 1DFJ | 81.43 | 97.85 | 95.87 | 0.80 |
| 1BUH | 100.00 | 99.06 | 99.15 | 0.95 |
| *Medium* |  |  |  |  |
| 1ACB | 82.35 | 98.87 | 97.00 | 0.85 |
| 1IB1 | 77.46 | 97.12 | 94.89 | 0.75 |
| 1IJK | 87.69 | 98.68 | 97.08 | 0.88 |
| 2NZ8 | 78.36 | 95.96 | 93.57 | 0.77 |
| *Difficult* |  |  |  |  |
| 1JZD | 74.33 | 97.51 | 94.83 | 0.75 |
| 1PXV | 93.70 | 86.39 | 87.19 | 0.85 |
| 2C0L | 68.25 | 99.11 | 94.26 | 0.77 |
| 2O3B | 82.35 | 95.87 | 97.00 | 0.82 |
| 2OT3 | 73.27 | 86.15 | 89.02 | 0.71 |

Results from Table 3 show that MCC values of 75% of test complexes are more than 0.80, and conclude a strong prediction. MCC values of other 4 test complexes are greater than 0.75 and less than 0.80 can be concluded as good in prediction. But, these complexes have more deviation in RMS values.

#### 3.2.3 Comparison with popular Servers

RMSD values of the best-docked structures obtained from Cons_SaDE are compared with the results of five popular docking web servers such as ZDOCK (Pierce et al. 2014), GRAMM-X (Tovchigrechko and Vakser 2006), ClusPro (Kozakov et al. 2013), Patch Dock (Schneidman-Duhovny et al. 2005) and Hex (Macindoe et al. 2010). The input, Protein 1 and Protein 2 of each test complex, is fed into the servers. The top-ranked docked structures from the servers are considered for RMSD calculations. Table 4 shows RMSD values of Cons_SaDE and other web servers.

**Table 4:**
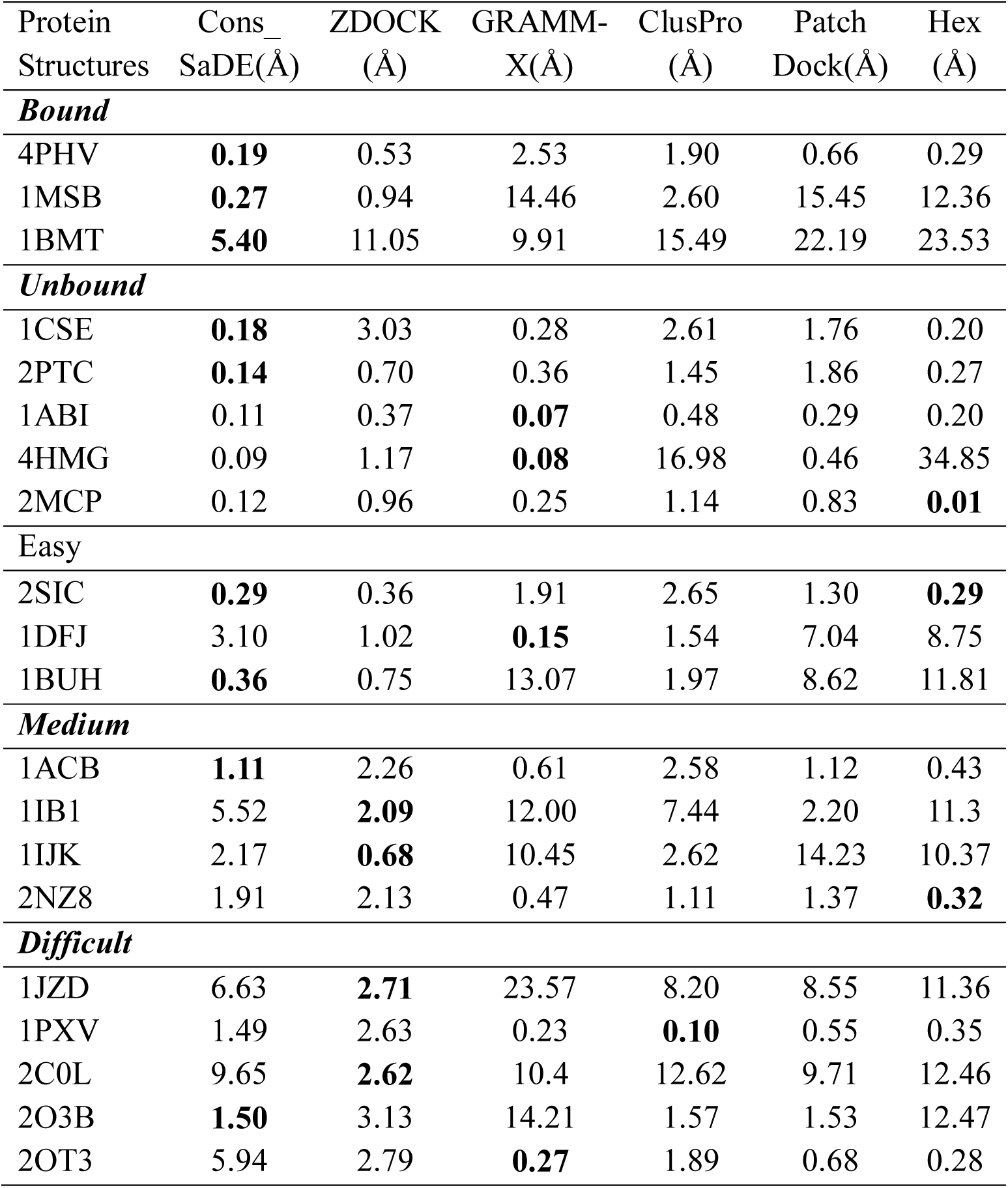
Comparison of RMSD values

From the results of Table 4, it is observed that among 20 test complexes, Cons_SaDE is producing good RMSD values for 9 complexes (4PHV, 1MSB, 1BMT, 1CSE, 2PTC, 2SIC, 1BUH, 1ACB, and 2O3B), whereas ZDOCK produces good RMSD values for 4 complexes (1IB1, 1IJK, 1JZD and 2C0L), GRAMM-X for 4 complexes (1ABI, 4HMG, 1DFJ and 2OT3), ClusPro for 1 complex (1PXV), and Hex for 3 complexes (2MCP, 2SIC and 2NZ8). The observation is that Cons_SaDE is reliable in producing good results for bound, unbound and easy complexes, whereas it is able to produce better RMSD values for medium and difficult categories among all servers.

To show the performance reliability of Cons_SaDE, RMSD values of top 10 models from Cons_SaDE and other servers are considered and averaged. The average RMSD values of top 10 models in Table 5 results that Cons_SaDE produces lower average RMSD values than all other servers. It is concluded that the top 10 models of Cons_SaDE are closer to each other and are not random like in other servers.

**Table 5:** Comparison of Average RMSD values

| Protein Structures | Cons_SaDE(Å) | ZDOCK (Å) | GRAMM-X(Å) | ClusPro (Å) | Patch Dock(Å) | Hex (Å) |
| --- | --- | --- | --- | --- | --- | --- |
| ***Bound*** |  |  |  |  |  |  |
| 4PHV | 0.36 | 4.45 | 11.96 | 7.54 | 4.02 | 3.84 |
| 1MSB | 0.20 | 7.30 | 17.68 | 10.42 | 16.91 | 16.70 |
| 1BMT | 7.03 | 21.63 | 20.24 | 22.12 | 23.79 | 24.15 |
| ***Unbound*** |  |  |  |  |  |  |
| 1CSE | 0.44 | 13.76 | 8.63 | 9.64 | 10.43 | 7.60 |
| 2PTC | 0.23 | 6.40 | 10.28 | 6.71 | 8.48 | 6.96 |
| 1ABI | 0.16 | 2.41 | 10.87 | 9.17 | 5.89 | 0.51 |
| 4HMG | 0.49 | 27.31 | 36.78 | 21.14 | 9.47 | 35.55 |
| 2MCP | 1.03 | 17.58 | 19.98 | 18.77 | 17.09 | 17.92 |
| ***Easy*** |  |  |  |  |  |  |
| 2SIC | 0.31 | 5.83 | 5.28 | 8.98 | 13.52 | 13.92 |
| 1DFJ | 7.41 | 5.26 | 13.75 | 6.41 | 13.87 | 12.48 |
| 1BUH | 0.58 | 9.99 | 18.39 | 10.98 | 11.81 | 13.32 |
| ***Medium*** |  |  |  |  |  |  |
| 1ACB | 0.23 | 5.73 | 8.57 | 6.03 | 15.51 | 11.50 |
| 1IB1 | 9.23 | 11.85 | 15.91 | 13.42 | 13.95 | 16.60 |
| 1IJK | 7.62 | 8.87 | 17.56 | 12.24 | 17.16 | 17.25 |
| 2NZ8 | 8.41 | 16.74 | 15.62 | 16.89 | 14.84 | 14.52 |
| ***Difficult*** |  |  |  |  |  |  |
| 1JZD | 10.69 | 11.68 | 24.64 | 12.45 | 10.88 | 12.72 |
| 1PXV | 5.40 | 9.20 | 14.43 | 13.56 | 14.07 | 9.18 |
| 2C0L | 10.46 | 18.24 | 13.49 | 19.94 | 17.42 | 20.21 |
| 2O3B | 8.43 | 10.27 | 20.42 | 13.01 | 15.13 | 17.31 |
| 2OT3 | 8.63 | 3.23 | 12.95 | 8.01 | 12.30 | 15.73 |

From Table 4, it is observed that few servers produce good RMSD values for certain complexes. For example, ZDOCK performs well in medium and difficult categories, GRAMM-X, Hex and Patch Dock servers do well in unbound category. While observing the results, the servers are not consistently producing good results for all the categories. In order to compare the performance of Cons_SaDE with the servers, ranking based nonparametric statistical tests are performed.

#### 3.2.4 Ranking based on nonparametric statistical tests

Here, nonparametric statistical tests of multiple comparisons (1xN) with control method (Derrac et al. 2011) are implemented to illustrate the performance of Cons_SaDE. The RMSD values from Table 4 are considered for tests. Table 6 shows the results of rankings of various algorithms using Friedman, Friedman Aligned, and Quade tests. From the rankings of algorithms, it is known that Cons_SaDE receives the lowest rank of 2.4, 41.6, and 2.1 for the Friedman, Friedman Aligned, and Quade tests respectively, by highlighting this as the best performing algorithm of comparison.

##### Performance comparison

Cons_SaDE is ranked first. The performances of the other methods are evaluated by taking Cons_SaDE as a control method. The hypotheses of equality between Cons_SaDE and others can be tested by the application of a set of post-hoc procedures. This post-hoc test can lead obtaining a p-value which determines the degree of rejection of each hypothesis. The level of significance is α =0.05. Tables 7- 9 show the adjusted p-values obtained using Friedman, Friedman Aligned and Quade tests, respectively and the p-values conclude the tests in the following manner:

**Table 6:** Average Ranking of the Algorithms

| ALGORITHM | AVERAGE RANKINGS |  |  |
| --- | --- | --- | --- |
|  | FRIEDMAN | ALIGNED<br>FRIEDMAN | QUADE |
| Cons_SaDE | 2.4 | 41.6 | 2.1 |
| ZDOCK | 3.35 | 44.95 | 2.8 |
| GRAMM-X | 3.48 | 65.33 | 3.7 |
| ClusPro | 4.14 | 63.55 | 3.8 |
| Patch Dock | 4.1 | 71.65 | 4.1 |
| Hex | 3.53 | 75.93 | 4.1 |

From Table 7, Friedman test shows a significant improvement of Cons_SaDE over ClusPro and Patch Dock for all the post-hoc procedures considered. The Li test exhibits the most powerful behavior, reaching the lowest p-values in comparisons. The adjusted p-values from *PBonf* to *PRom* in Hex and GRAMM-X are more than significant level implies that they are as good as Cons_SaDE with little deficiencies. ZDOCK server challenges Cons_SaDE results.

**Table 7:**
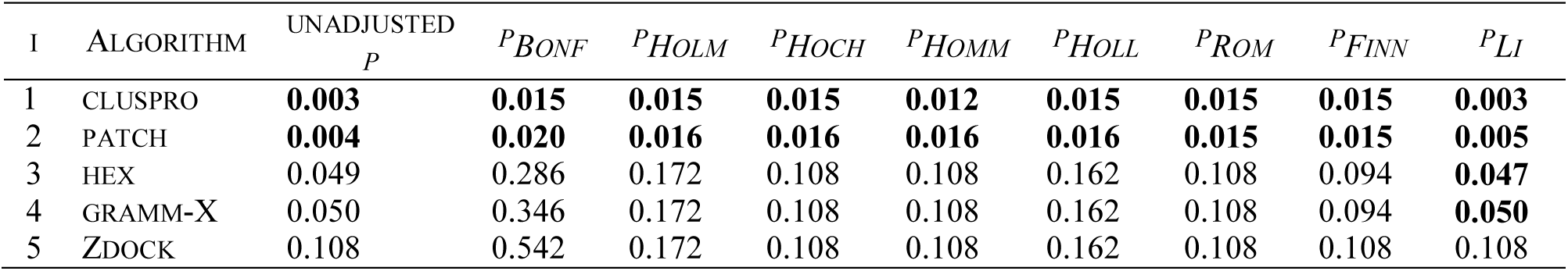
Adjusted p-values for the Friedman Test

According to the results of Aligned Friedman test from Table 8, Cons_SaDE exhibits better performance than Hex and Patch Dock servers.

**Table 8:**
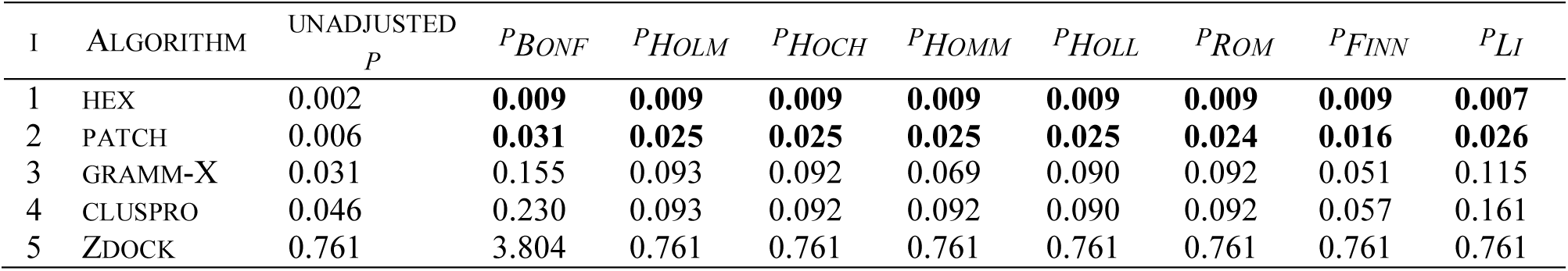
Adjusted p-values for the Aligned Friedman Test

The Quade test results, as shown in Table 9, does not find any significant performance difference between Cons_SaDE and other algorithms. In order to make this algorithm competitive in all cases, a new constraint may be introduced i.e. the nature of interaction between proteins can be considered as one of the optimization constraints.

**Table 9:** Adjusted p-values for the Quade Test

| I | ALGORITHM | UNADJUSTED<br>$P$ | $P_{BONF}$ | $P_{HOLM}$ | $P_{HOCH}$ | $P_{HOMM}$ | $P_{HOLL}$ | $P_{ROM}$ | $P_{FINN}$ | $P_{LI}$ |
| --- | --- | --- | --- | --- | --- | --- | --- | --- | --- | --- |
| 1 | PATCH | 0.057 | 0.287 | 0.287 | 0.255 | 0.178 | 0.256 | 0.243 | 0.256 | 0.101 |
| 2 | HEX | 0.064 | 0.319 | 0.287 | 0.255 | 0.191 | 0.256 | 0.243 | 0.256 | 0.111 |
| 3 | CLUSPRO | 0.108 | 0.541 | 0.325 | 0.266 | 0.216 | 0.291 | 0.266 | 0.256 | 0.176 |
| 4 | GRAMM-X | 0.133 | 0.666 | 0.325 | 0.266 | 0.266 | 0.291 | 0.266 | 0.256 | 0.208 |
| 5 | ZDOCK | 0.492 | 2.460 | 0.492 | 0.492 | 0.492 | 0.492 | 0.492 | 0.492 | 0.492 |

As an outcome, this study has provided the following significant observations.

i. Introduction of these constraints made a guided search for the better/best promising solutions for many complexes.
ii. From the results of Quade test, it is observed that the nature of interactions needs to be probably taken into consideration as a constraint in Cons_SaDE to make this algorithm competitive in all cases.
iii. Also, all 176 complexes from Benchmark 4.0 need to be implemented as an extended work to show the competitive performance of Cons_SaDE.

## 4. Conclusion

The main components of protein docking are effectively handled in this work. Modified rigid docking representation, called Rotate Only Representation, is proposed to improve the search efficiency by producing more number of promising solutions with minimum energy. The computational cost of searching for the optimal solutions from the huge search space can be reduced using guided search. The guided search is achieved by introducing three constraints, such as boundary surface complementary interactions, non-bonded inter-atomic allowed distances, and finding of interaction and non-interaction sites, into this docking problem. Now, the problem is formulated as a constrained optimization problem with an objective of minimizing potential energy, satisfying the constraints. Cons_SaDE is used as a search algorithm, which efficiently explores the promising solutions. To test the performance of this algorithm, few bound and unbound complexes from Protein Data Bank (PDB) and few easy, medium difficult and difficult complexes from Zlab benchmark 4.0 set are used. From the results of potential energy, buried surface area, RMSD, Matthew’s Correlation Coefficient etc. for the considered complexes prove that Cons_SaDE is able to produce improved solutions consistently.

RMSD values of the best conformations obtained from five popular docking web servers are compared with Cons_SaDE results, and nonparametric statistical tests of multiple comparisons (1xN), with control method, are implemented to show the relative performance of this proposed algorithm. Cons_SaDE is successful in generating low RMSD structures with better sensitivity, specificity, accuracy and correlation coefficient measures across all categories of data sets except easy category. From the results of various metrics and nonparametric statistical tests, it is concluded that Cons_SaDE has produced good quality of solutions for most of the data sets considered. Still to improve the performance of Cons_SaDE, it is necessary to consider the size and nature of interactions of proteins as constraints.

## ACKNOWLEDGMENT

The first author takes this opportunity to express her profound gratitude and deep regards to Ms. P.J. Eswari Pandaranayaka, Postdoctoral Research Scholar, MKU, for her exemplary support by providing valuable information and guidance and constructive feedback on the evaluation of the results of this work. The first author is obliged to Mrs. C.V. Nisha Angeline, Asst. Prof, I.T, for her assistance in initial coding. The first author is thankful to Mr. G.Vivek, Software Engineer, Eicsson, for his constant support by means of facilitating the cluster installation and debugging, essential for this work.

## References

1. Bajaj C, Chowdhury R, Siddavanahalli V (2011) F2Dock: Fast Fourier Protein-Protein Docking. IEEE/ACM Transactions on Computational Biology and Bioinformatics 8(1):45-58

2. Banting L, Clark T, Thurston DE (2012) Drug Design Strategies: Computational Techniques and Applications 1 edition. Royal Society of Chemistry, London, UK, 2012.

3. Baxter CA, Murray CW, Clark DE, Westhead DR, Eldridge MD (1998) Flexible docking using Tabu search and an empirical estimate of binding affinity. Proteins 33(3): 367-382

4. Cai Y, Wang J, Yin J (2012) Learning-enhanced differential evolution for numerical optimization. Soft Computing 16(2): 303-330

5. Cai X, Hu Z, Fan Z (2013) A novel memetic algorithm based on invasive weed optimization and differential evolution for constrained optimization. Soft Computing, 17(10): 1893-1910

6. Chaudhury S, Gray JJ (2008) Conformer selection and induced fit in flexible backbone protein protein docking using computational and NMR ensembles. J Mol Biol 2008; 381(4): 1068-1087. doi:10.1016/j.jmb.2008.05.042

7. Chen K, Li T, Cao T (2006) Tribe-PSO: A novel global optimization algorithm and its application in molecular docking. Journal of Chemometrics and Intelligent Laboratory Systems 82(1): 248-259

8. Chen R, Li L,Weng Z (2003) Zdock: An initial-stage protein-docking algorithm, Proteins: Structure, Function, and Genetics, Special Issue: CAPRI - Critical Assessment of PRedicted Interactions. Proteins 52(1):80–87

9. Clark KP (1995) Flexible ligand docking without parameter adjustment across four ligand-receptor complexes. Journal of Computational Chemistry 16(10): 1210-1226

10. Coello CAC, Montes EM (2002) Constraint-handling in genetic algorithms through the use of dominance-based tournament selection. Advanced Engineering Informatics, 16(3): 193-203

11. Correlation (2016), https://en.wikipedia.org/wiki/Matthews_correlation_coefficient

12. de Vries S, Zacharias M (2013) Flexible docking and refinement with a coarse-grained protein model using ATTRACT. Proteins 81(12) 2167–2174

13. de Vries SJ, van Dijk M, Bonvin AM (2010) The HADDOCK web server for data-driven biomolecular docking. Nature Protocols, 5: 883–897

14. Derrac J, García S, Molina D, Herrera F (2011) A practical tutorial on the use of nonparametric statistical tests as a methodology for comparing evolutionary and swarm intelligence algorithms. Swarm and Evolutionary Computation 1(1) 3-18

15. dssp, (2012), Centre for Molecular and Biomolecular Informatics. http://swift.cmbi.ru.nl/gv/dssp

16. Esquivel-Rodrigue J, Yang YD, Kihara D (2012) Multi-LZerD: multiple protein docking for asymmetric complexes. Proteins 80(7): 1818–1833

17. Esquivel-Rodríguez J, Kihara D (2012) Effect of conformation sampling strategies in genetic algorithm for multiple protein docking BMC Proc. 6 (Suppl 7): S4

18. Fernandez-Recio J, Totrov M, Abagyan R (2003) ICM-DISCO docking by global energy optimization with fully flexible side-chains. Proteins 52(1) 113–117

19. Gabb HA, Jackson RM, Sternberg MJ (1997) Modelling protein docking using shape complementarity, electrostatics and biochemical information. Journal of Molecular Biology 272(1): 106–120

20. Garzon JI, Lopéz-Blanco JR, Pons C, Kovacs J, Abagyan R, Fernandez-Recio J, Chacon P (2009) FRODOCK: a new approach for fast rotational protein–protein docking. Bioinformatics 25(9): 2544–255

21. Gray JJ, Moughan SE, Wang C, Schueler-Furman O, Kuhlman B, Rohl CA, Baker D (2003) Protein-Protein Docking with Simultaneous Optimization of Rigid-Body Displacement and Side-Chain Conformations. Journal of Molecular Biology 331(1): 281-299

22. Hashmi I, Shehu A (2012) HopDock: a probabilistic search algorithm for decoy sampling in protein-protein docking. Proteome Science 11 Supplement 1

23. Huang P, Love JJ, Mayo SL (2005) Adaptation of a Fast Fourier Transform-Based Docking Algorithm for Protein Design. Journal of Computational Chemistry 26(12): 1222–1232

24. Jones G, Willett P, Glen RC, Leach AR, Taylor R (1997) Development and validation of a genetic algorithm for flexible docking. Journal of Molecular Biology 267(3): 727-748

25. Kong X, Ouyang H, Piao X (2013) A prediction-based adaptive grouping differential evolution algorithm for constrained numerical optimization. Soft Computing, 17(12): 2293-2309

26. Korb O, Stutzle T, Exner TE (2006) PLANTS: Application of ant colony optimization to structure-based drug design. In Proceedings of Ant Colony Optimization and Swarm Intelligence, 5th International Workshop pp. 247-258

27. Kozakov D, Beglov D, Bohnuud T, Mottarella S, Xia B, Hall DR, Vajda S (2013) How good is automated protein docking? Proteins: Structure, Function, and Bioinformatics 81(12): 2159-2166

28. Kozakov D, Brenke R, Comeau SR, Vajda S (2006) PIPER: An FFT-based protein docking program with pairwise potentials. Proteins 65(2):392-406

29. Kuntz ID, Blaney JM, Oatley SJ, Langridge R, Ferrin TE (1982) A geometric approach to macromolecule–ligand interactions. Journal of Molecular Biology 161(2) 269–288

30. Li B, Kihara D (2012) Protein docking prediction using predicted protein-protein interface. BMC Bioinformatics 13(7):1-17

31. Li L, Guo D, Huang Y, Liu S, Xiao Y (2011) ASPDock: protein-protein docking algorithm using atomic solvation parameters model. BMC Bioinformatics 12(36)

32. Macindoe G, Mavridis L, Venkatraman V, Devignes MD, Ritchie DW (2010) HexServer: an FFT-based protein docking server powered by graphics processors. Nucleic Acids Research 38: W445-W449

33. Mashiach E, Nussinov R, Wolfson HJ (2010) FiberDock: Flexible induced-fit backbone refinement in molecular docking. Proteins 78(6): 1503-19

34. Michalewicz Z, Schoenauer M (1996) Evolutionary algorithms for constrained parameter optimization problems. Evolutionary Computation, MIT Press Journal, 4(1): 1-32

35. Moal IH, Bates PA (2010) SwarmDock and the use of normal modes in protein–protein docking. International Journal of Molecular Science 1(10): 3623-3648

36. Morris GM, Goodsell DS, Halliday RS, Huey R, Hart WE, Belew RK, Olson AJ (1998) Automated docking using a Lamarckian genetic algorithm and empirical binding free energy function. Journal of Computational Chemistry 19(14): 1639-1662

37. Oostenbrink C, Villa A, Mark AE, Van Gunsteren WF (2004) A Biomolecular Force Field Based on the Free Enthalpy of Hydration and Solvation: The GROMOS Force-Field Parameter Sets 53A5 and 53A6. Weily Journal of Computational Chemistry 25(13): 1656–1676

38. Pei J, Wang Q, Liu Z, Li Q, Yang KL, Lai L (2006) PSI-DOCK: Towards highly efficient and accurate flexible ligand docking. Proteins 62(4): 934-946

39. Pierce BG, Wiehe K, Hwang H, Kim BH, Vreven T, Weng Z (2014) ZDOCK Server: Interactive Docking Prediction of Protein-Protein Complexes and Symmetric Multimers. Bioinformatics 30(12): 1771-1773

40. Protein Docking Benchmark – Zlab (2010), https://zlab.umassmed.edu/benchmark/

41. Pymol, (2000) http://pldserver1.biochem.queensu.ca/~rlc/work/teaching/BCHM823/pymol/alignment/

42. Qin AK, Huang VL, Suganthan PN (2009) Differential Evolution Algorithm With Strategy Adaptation for Global Numerical Optimization IEEE transactions on Evolutionary Computation 13(2): 398-417

43. Reid DJ (1996), Genetic algorithms in constrained optimization. Mathematical and Computer Modelling, Elsevier, 23(5): 87-111

44. Ritchie DW, Kozakov D, Vajda S (2008) Accelerating and focusing protein-protein docking correlations using multi-dimensional rotational FFT generating functions. Bioinformatics 24(17): 1865–187

45. Roberts VA, Thompson EE, Pique ME, Perez MS, Ten Eyck LF (2013) DOT2: macromolecular docking with improved biophysical models. Journal of Computational Chemistry 34(20): 1743–1758

46. Schneidman-Duhovny D, Inbar Y, Nussinov R, Wolfson HJ (2005) PatchDock and SymmDock: servers for rigid and symmetric docking Nucleic Acids Research 33: W363-367

47. Schneidman-Duhovny D, Inbar Y, Nussinov R, Wolfson HJ (2005) PatchDock and SymmDock: servers for rigid and symmetric docking. Nuclic Acids Research 33: W363-367

48. Storn R, Price KV (1997) Differential evolution-A simple and efficient heuristic for global Optimization over Continuous Spaces. Journal of Global Optimization 11(4): 341-359

49. Stroganov O V, Novikov FN, Stroylov VS, Kulkov V, Chilov GG (2008) Lead finder: An approach to improve accuracy of protein-ligand docking, binding energy estimation, and virtual screening. Journal of Chemical Information and Modeling 48(12): 2371-2385

50. Suenaga A, Okimoto N, Hirano Y, Fukui K (2012) An Efficient Computational Method for Calculating Ligand Binding Affinities. PLoS ONE 7(8): e42846. https://doi.org/10.1371/journal.pone.0042846.

51. Takahama T, Sakai S (2009) Solving Difficult Constrained Optimization Problems by the ε Constrained Differential Evolution with Gradient-Based Mutation. Constraint-Handling in Evolutionary Optimization, Springer 198: 51-72

52. Thomsen R, Christensen MH (2006) MolDock: A new technique for high accuracy molecular docking. Journal of Medicinal Chemistry 49(11): 3315-3321

53. Tovchigrechko A, Vakser IA (2006) GRAMM-X public web server for protein-protein docking. Nucleic Acids Research 34:W310-314

54. Tovchigrechko A,Vakser IA (2005) Development and testing of an automated approach to protein docking. Proteins 60(2):296-301

55. Wang C, Bradley P, Baker D (2007) Protein–Protein Docking with Backbone Flexibility. J Mol Biol. 373(2): 503-19

